# The DNA Methylation Landscape of Giant Viruses

**DOI:** 10.1101/2019.12.21.884833

**Authors:** Sandra Jeudy, Sofia Rigou, Jean-Marie Alempic, Jean-Michel Claverie, Chantal Abergel, Matthieu Legendre

## Abstract

DNA methylation is an important epigenetic mark that contributes to various regulations in all domains of life. Prokaryotes use it through Restriction-Modification (R-M) systems as a host-defense mechanism against viruses. The recently discovered giant viruses are widespread dsDNA viruses infecting eukaryotes with gene contents overlapping the cellular world. While they are predicted to encode DNA methyltransferases (MTases), virtually nothing is known about the DNA methylation status of their genomes. Using single-molecule real-time sequencing we studied the complete methylome of a large spectrum of families: the Marseilleviridae, the Pandoraviruses, the Molliviruses, the Mimiviridae along with their associated virophages and transpoviron, the Pithoviruses and the Cedratviruses (of which we report a new strain). Here we show that DNA methylation is widespread in giant viruses although unevenly distributed. We then identified the corresponding viral MTases, all of which are of bacterial origins and subject to intricate gene transfers between bacteria, viruses and their eukaryotic host. If some viral MTases undergo pseudogenization, most are conserved, functional and under purifying selection, suggesting that they increase the viruses’ fitness. While the Marseilleviridae, Pithoviruses and Cedratviruses DNA MTases catalyze N^6^-methyl-adenine modifications, some MTases of Molliviruses and Pandoraviruses unexpectedly catalyze the formation of N^4^-methyl-cytosine modifications. In Marseilleviridae, encoded MTases are paired with cognate restriction endonucleases (REases) forming complete R-M systems. Our data suggest that giant viruses MTases could be involved in different kind of virus-virus interactions during coinfections.

## Introduction

Methylation of DNA is an important class of epigenetic modification observed in the genomes of all domains of life. In eukaryotes, it is involved in biological processes as diverse as gene expression regulation, transposon silencing, genomic imprinting or development (1–4). In prokaryotes, DNA methylation often results from the targeted activity of MTases that are components of the R-M systems, which involve methylation and restriction activity. Within these systems, REases cleave the DNA only if the shared recognized motifs are unmethylated (5). This provides prokaryotes with a powerful weapon against foreign DNA, such as the one of infecting viruses (6). Besides R-M systems, prokaryotic DNA MTases may occur without cognate REases, in which case they are coined “orphan”. Prokaryotic orphan MTases are involved in the regulation of gene expression (7, 8), DNA replication (9), DNA repair (10) and cell cycle regulation (11).

Outside of the cellular world, some DNA viruses exploit DNA methylation as a mechanism to regulate their replication cycle. For instance, the transition from latent to lytic infection in Epstein-Barr Virus is mediated by the expression of genes that are silenced or transcribed according to the methylation status of their promoter (12, 13). Iridoviruses and ascoviruses sometimes exhibit heavily methylated genomes and encode their own MTases (12, 14). Phycodnaviridae also encode functional MTases able to methylate their own DNA (15–17), and among them chloroviruses encode complete R-M systems with their associated REases (18, 19). These endonucleases, packaged in the virions, contribute to the degradation of host DNA either to allow for the recycling of deoxynucleotides, or to inhibit the expression of host genes by shifting the transcription from host to viral DNA (19).

Over the last fifteen years several viruses whose particles are large enough to be seen by light microscopy were discovered (20–30). These so-called “giant viruses” exhibit DNA genomes as large and complex as prokaryotes (21), or even parasitic eukaryotes (29). A growing body of metagenomics surveys shows that they are widespread on the planet in all kinds of environments (31–36). The first family of giant viruses to be discovered, the Mimiviridae, have megabase-sized AT-rich linear genomes packaged in icosahedral capsids (21–23). In contrast, the Pandoraviruses have GC-rich linear genomes, twice as big with up to 2.5 Mb, packaged in amphora-shaped capsids (29, 37). Again different, Pithoviruses (26) and Cedratviruses (27, 38) have smaller circular AT-rich genomes, ranging from 400 to 700 Mb, and packaged in the largest known amphora-shaped capsids. Thus, although these different giant viruses infect the same hosts (amoebas of the acanthamoeba genus), they exhibit different morphological features, replication cycles, gene contents, and potential epigenomic modifications. To date virtually nothing is known about the epigenomes of these giant viruses. In particular, the methylation status of their DNA is unknown despite the presence of predicted MTases in their genomes. Pandoravirus dulcis for instance encodes up to five different DNA MTases (UniprotKB IDs: S4VR68, S4VTY0, S4VS49, S4VUD3 and S4VQ82) (29). Yet, it remains to be assessed whether any of these enzymes methylate the viral DNA.

Most of the genome-wide studies of eukaryotic DNA methylation have been performed using bisulfite sequencing techniques (39). These approaches only detect 5-methyl-cytosine modifications and are thus not well suited for the analysis of prokaryotic-like epigenomic modifications, mostly composed of N^6^-methyl-adenines and N^4^-methyl-cytosines. However, the recently developed single-molecule real-time (SMRT) sequencing method overcomes this limitation (40, 41). Briefly this approach analyses the kinetics of incorporation of modified nucleotides by the polymerase compared to the non-modified ones. The Inter-Pulse Duration ratio (IPDr) metric can then be computed for each genomic position, and makes it possible to map all modified nucleotides and methylated motifs along the genome. This approach is now extensively used to study the methylation landscapes of isolated bacteria (42), archaea (43) and even prokaryotic metagenomes (44).

Here we used SMRT sequencing to survey the complete methylome of a large spectrum of giant viruses. We analyzed two distinct Mimiviridae and their associated transpovirons as well as a virophage (45). We also surveyed a Marseilleviridae (46), five distinct Pandoraviruses (29, 37, 47), a Mollivirus (48) and a Pithovirus (26). Finally, we isolated a new Cedratvirus (Cedratvirus kamchatka) that we sequenced using SMRT sequencing to assess its methylome. Furthermore, we thoroughly annotated MTases and REases contained in all these genomes and analyzed their phylogenetic histories. Our findings reveal that DNA methylation is widespread among giant viruses and open new avenues of research on its role in their population dynamics.

## Results

### Methylome and MTase gene contents of giant viruses

We gathered available PacBio SMRT sequence data from previously published genomes of diverse families to analyze the DNA methylation profile of a wide of range of giant viruses. SMRT genomic data were collected for the following viruses: two *Mimiviridae* members (*Megavirus vitis* and *Moumouvirus australiensis*), one *Lavidaviridae* member (*Zamilon vitis*), two transpovirons (*Moumouvirus australiensis* transpoviron and *Megavirus vitis* transpoviron), one *Marseilleviridae* member (Melbournevirus) and five Pandoraviruses (*Pandoravirus celtis*, *Pandoravirus dulcis*, *Pandoravirus neocaledonia*, *Pandoravirus quercus* and *Pandoravirus salinus*). The used datasets are listed in Table S1A. In addition, we resequenced the complete genomes of *Mollivirus sibericum* and *Pithovirus sibericum* together with a newly isolated strain of Cedratvirus (*Cedratvirus kamchatka*) on the PacBio platform. The sequence data obtained from the whole collection corresponds to an average coverage of 192-fold.

We then aligned the SMRT reads to their corresponding reference genomes (see Table S1A) and computed IPDr at each genomic position (see Methods). These genome-wide profiles were used to identify overrepresented sequence motifs at positions with high IPDr values. All the identified motifs were palindromic and prone to either N^4^-methyl-cytosine or N^6^-methyl-adenine methylations (Fig. 1). As a control, we applied the same procedure to DNA samples of *Mollivirus sibericum* and *Pithovirus sibericum* subjected to whole genome amplifications (WGA), which in principle erase methylation marks (41). As expected, no overrepresented methylated motif was detected in these controls, and the median IPDr of the motifs previously detected in the wild-type datasets were basal here (see Fig. S1 and Fig. 1).

**Figure 1.**
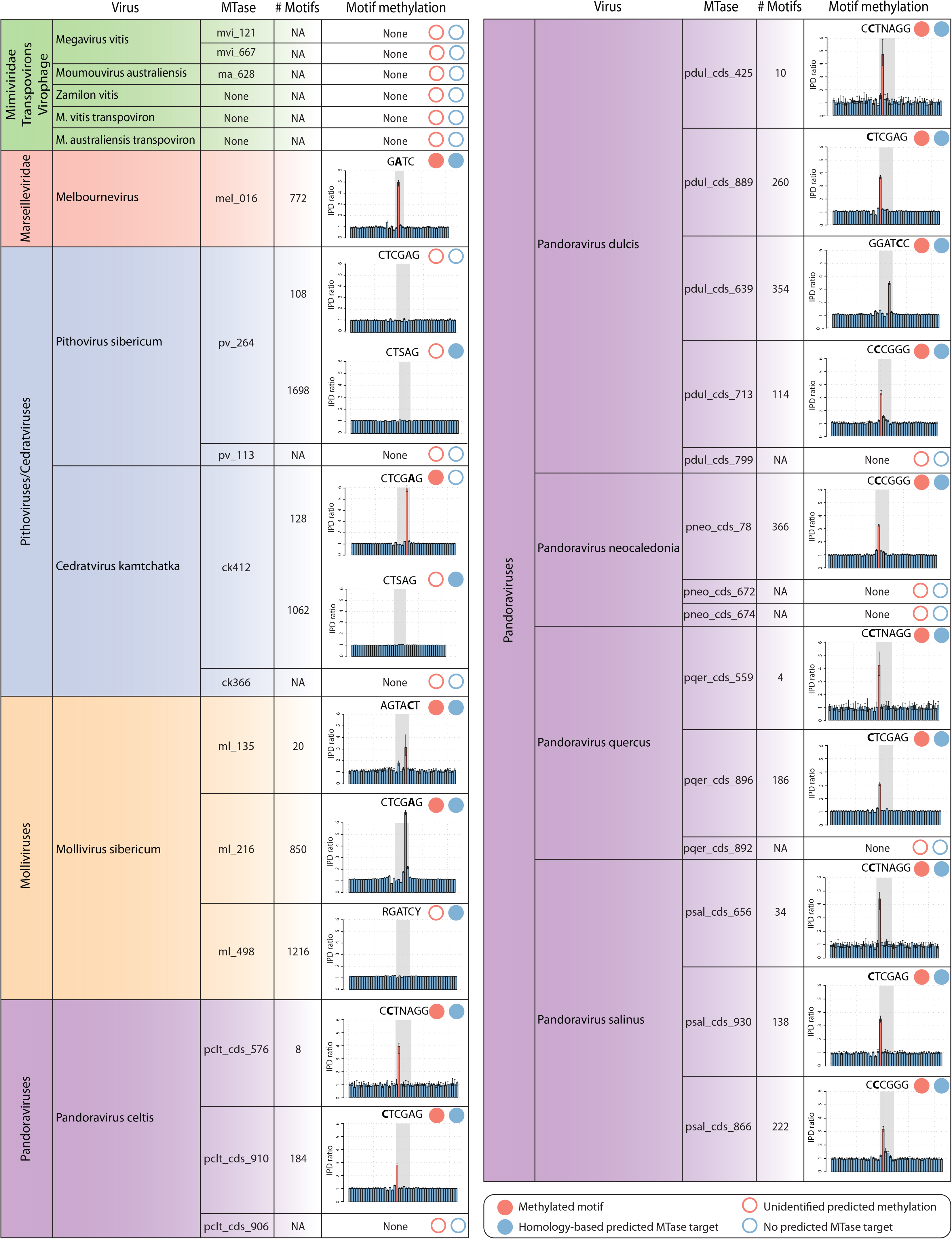
Encoded MTases and targeted methylated motifs in the giant viruses’ genomes. The encoded DNA MTases of each virus are shown along with the number of occurrences of the predicted targets (if any) on both strands of the cognate genomic sequence. Experimentally verified from SMRT data methylated motifs (filled circle) or unidentified predicted methylation (empty circle) are depicted using red circles. Likewise, predicted (filled circle) and not predicted (empty circle) targets based on sequence homology of the encoded MTase are shown using blue circles. Methylated nucleotides are indicated using bold characters. The median IPDr profiles of the motifs (highlighted in gray) and their surrounding 20 nucleotides on each side are shown. Red bars correspond to positions with significantly high IPDr values. Error bars correspond to 95% intervals based on bootstrapping.

In parallel, we analyzed the DNA MTases encoded in these genomes and predicted their target sequences based on their homology with characterized MTases (see Methods). All the MTases for which a target site could be predicted were putative type II MTases. In most cases (78%) we found an agreement between the predicted targets and the detected methylated motifs (Fig. 1).

### *Marseilleviridae* members encode complete R-M systems

Melbournevirus encodes a DNA MTase (mel_016) predicted to target GATC sites. Our data confirm that G**A**TC motifs were modified (the methylated position is indicated in bold) with N^6^-methyl-adenines (Fig. 1). We then searched for the possible cognate REase in the genomic vicinity of mel_016 and identified the neighboring mel_015 gene as a candidate. Although the encoded protein does not exhibit a recognizable motif using standard domain search tools (49, 50), a search against Rebase, the database dedicated to R-M systems (51), identified it as a probable GATC-targeting REase. Moreover, the mel_015 protein is similar (blast E-value = 2×10^−33^) to the *Paramecium bursaria* Chlorella virus 1 (PBCV-1) CviAI REase known to target GATC sites. Melbournevirus thus encodes a complete R-M system.

The N^6^-methyl-adenine modification is typical of prokaryotic MTases. Since Melbournevirus is a eukaryotic virus, our finding immediately questioned the evolutionary history of its encoded R-M system. We thus reconstructed the phylogeny of the complete system, including the MTase and the REase (see Methods). The mel_016 MTase strikingly branches within the prokaryotes along with other viruses, mostly Chloroviruses and members of the *Marseilleviridae* and *Mimiviridae* of the proposed mesomimivirinae subfamily (52) (Fig. S2A). In agreement with its enzymatic activity, this phylogeny suggests that the Melbournevirus MTase was acquired from a prokaryote. Likewise, the phylogeny of the mel_015 REase suggests its prokaryotic origin (Fig. S2B). Altogether, these results support a relatively ancient acquisition from prokaryotes as the origin of the complete Marseilleviridae R-M system.

Surprisingly, we did not identify orthologues of the Melbournevirus R-M system in all *Marseilleviridae* members. As shown in Figure 2, only 5 out of the 13 Marseilleviruses genomes contain both a MTase and a REase, always encoded next to each other. The other marseilleviruses encode neither the MTase nor the REase. The *Marseilleviridae* phylogeny based on core genes (see Methods) clearly coincides with its dichotomous distribution (Fig. 2). All the clade A members of the *Marseilleviridae* encode a complete R-M system, while the others do not. This suggests that the Marseilleviridae R-M system was acquired by the clade A ancestor. It is worth noticing that once acquired, the R-M system was maintained, with none of the two enzymes undergoing pseudogenization. This suggests that the encoded R-M system has a functional role in these viruses.

**Figure 2.**
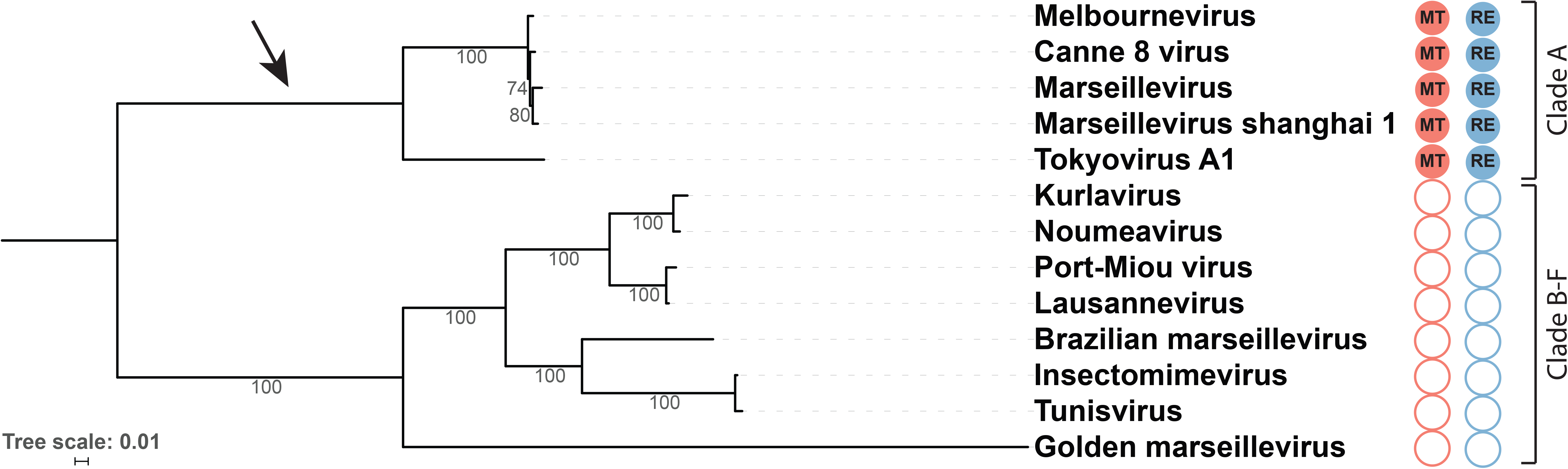
Phylogenetic reconstruction of the Marseilleviridae R-M system. Phylogeny of the Marseilleviridae completely sequenced viruses (see GenBank accessions in Table S1B) based on protein sequence alignments of the 115 strictly conserved single copy orthologues. The tree was calculated using the best model of each partitioned alignment as determined by IQtree (107). The red and blue filled circles highlight the presence of encoded Marseilleviridae R-M systems MTases and REases respectively. The empty circles highlight the absence of the MTase (in red) and the REase (in blue). The arrow points to the most parsimonious acquisition of an R-M system in the Marseilleviridae.

### Activity of the Marseilleviridae R-M system on non-self DNA

Once methylated by the mel_016-encoded enzyme, we expect the viral DNA to be protected from its digestion by the cognate mel_015 REase. If not, the Melbournevirus genome would be theoretically fragmented into 387 fragments of 954 nt on average. We verified this prediction by conducting DNA restriction experiments using two endonucleases targeting GATC sites: DpnI and DpnII. The former only cleaves DNA at modified G**A**TC sites containing a N^6^-methyl-adenine, while the latter conversely only cleaves unmethylated GATC sites. Figure 3 demonstrates that Melbournevirus DNA is digested by DpnI but not by DpnII. Thus, Melbournevirus protects its own genome from its encoded R-M system REase. As a control, we reproduced the above experiment with the DNA of Noumeavirus (53), a *Marseilleviridae* member belonging to the clade B that do not encode an R-M system (Fig. 2). As expected, the Noumeavirus genome is digested by DpnII but not by DpnI (Fig. 3). This demonstrates that Noumeavirus DNA is not methylated at G**A**TC sites and is thus susceptible to degradation by a co-infecting marseillevirus bearing a functional R-M system.

**Figure 3.**
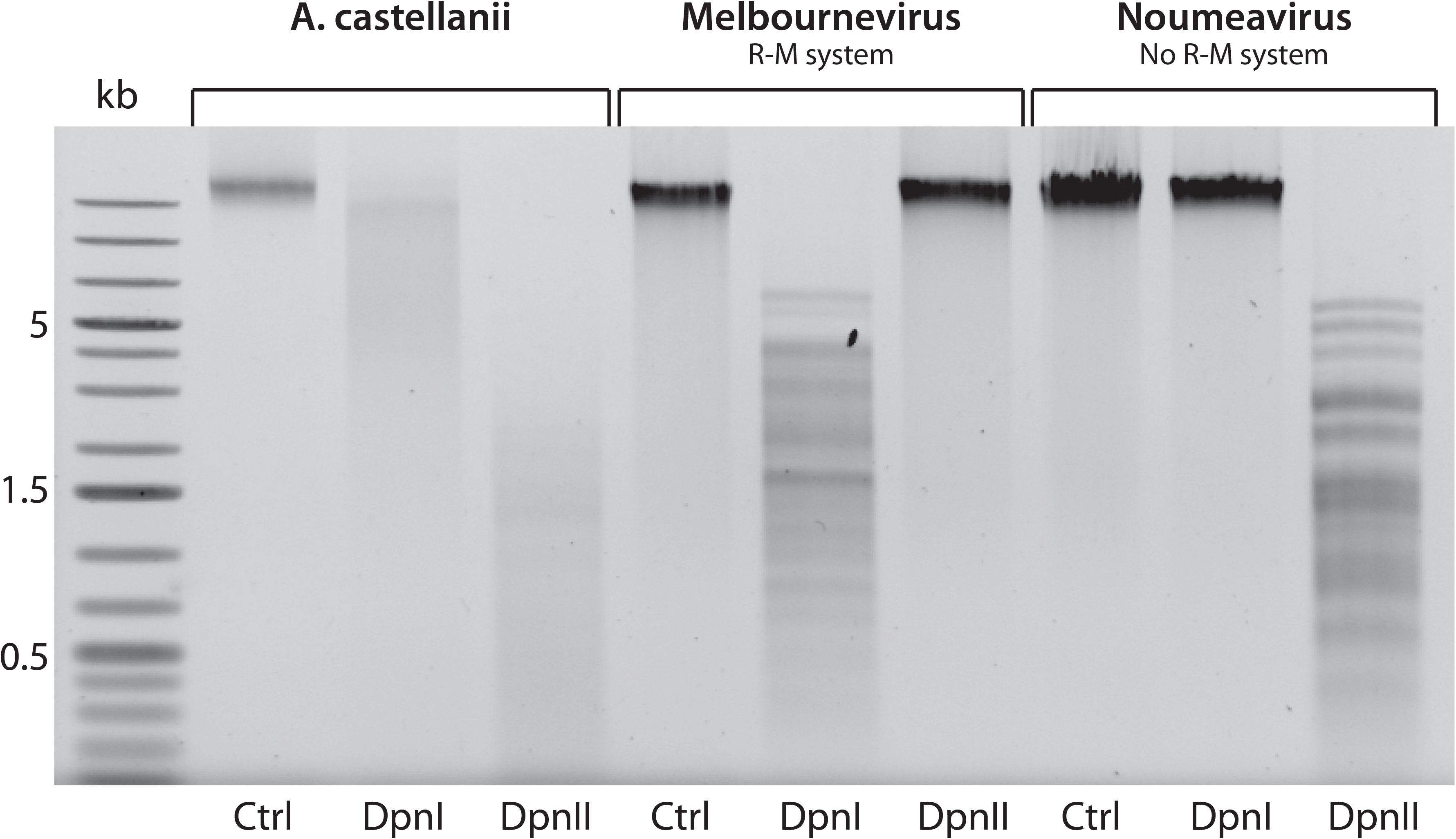
Host and Marseilleviridae DNA protection against GATC targeting REases. Agarose gel electrophoresis analysis of A. castellanii, Melbournevirus and Noumeavirus DNA digested with GATC targeting restriction enzymes. Restriction patterns using DpnI and DpnII enzymes are presented with control DNA. DpnI cleaves DNA at G**A**TC sites containing N^6^-methyl-adenines and DpnII at GATC sites containing unmethylated adenines.

To verify whether the *Acanthamoeba castellanii* host genome was sensitive to DNA degradation at GATC sites we performed digestion assays using the same enzymes. According to its sequence (AEYA01000001 to AEYA01002545 GenBank accessions), the *A. castellanii* genome should be fragmented into 171,298 pieces of 273 nt on average. Surprisingly, the *A. castellanii* genome is cleaved by both enzymes (Fig. 3). This indicates that the host DNA contains a mixture of methylated and unmethylated GATC sites. However, the restriction profiles show that unmethylated positions are in larger proportion than methylated ones.

Since the host DNA is (at least partially) unprotected from the Marseilleviridae encoded R-M system we next assessed its potential degradation during a Melbournevirus infection. As shown in Figure S3, *A. castellanii* DNA is not degraded during the infection. As expected, the infection of *A. castellanii* with Noumeavirus, which do not encode the GATC R-M system, do not alter its DNA. Hence, Marseilleviridae encoded R-M systems do not contribute to host DNA degradation.

To further investigate the role of the Marseilleviridae R-M systems we analyzed the timing of expression of both enzymes (the MTase and the REase) during a Melbournevirus infection. Figure 4 shows that the mel_015 REase gene is first transcribed between 30min and 45min post infection, followed by the mel_016 MTase gene expressed between 45min and 1h post infection. In addition, proteomic data of the Melbournevirus virion from (53) confirm that the mel_016 MTase protein is packaged in the particle whereas the REase is not.

**Figure 4.**
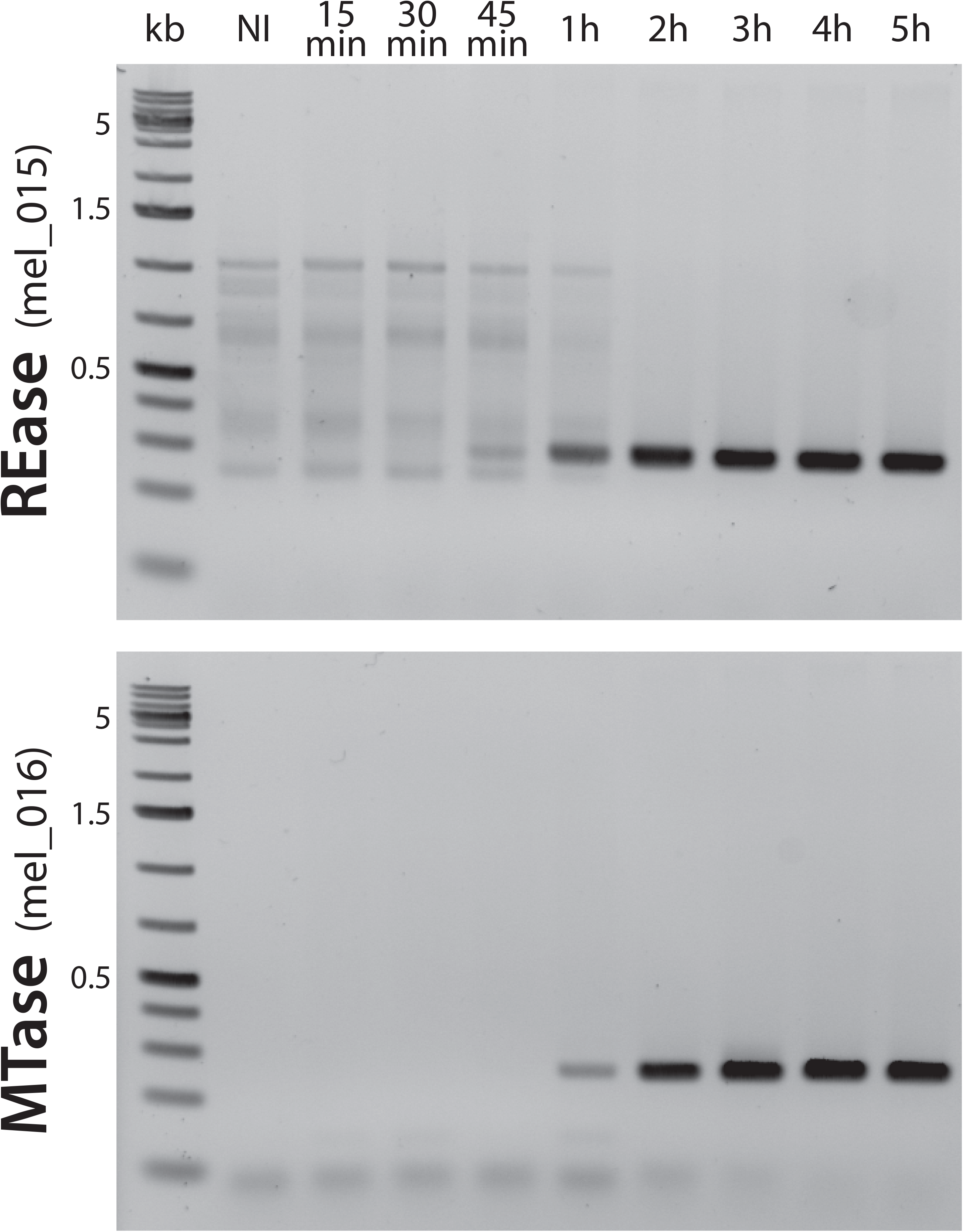
Expression timing of Melbournevirus R-M system MTase and REase. Shown are the RT-PCRs of the transcripts corresponding to the mel_015 and mel_016 genes during a melbournevirus infection. Times (post infection) are listed on the top of the figure. NI corresponds to non-infected.

### The newly isolated *Cedratvirus kamchatka*

Cedratviruses are giant viruses morphologically and to some extant genetically related to Pithoviruses (38). Four completely sequenced genomes are available today: *Cedratvirus* A11 (27), *Cedratvirus zaza* (54), *Cedratvirus lausannensis* (38) and *Brazilian Cedratvirus* (54). We recently isolated a new strain of Cedratvirus (named “*Cedratvirus kamchatka*”) from a muddy grit soil sample collected near a lake at kizimen volcano, Kamchatka (Russian Federation N 55°05′50 E 160°20′58) (see Methods). SMRT sequencing was used to characterize both its genome and methylome. The *C. kamchatka* genome was assembled into a circular 466,767 bp DNA molecule (41% G+C), predicted to encode 545 protein-coding genes The genome size and topology was confirmed by pulsed field gel electrophoresis (PFGE) (Fig. S4A).

The phylogenetic tree computed from the Pithoviruses and Cedratviruses core genes shows that they cluster in well-separated groups (Fig. S4B). Their orthologous proteins share an average of 46% identical residues. The available Cedratviruses appear to split into three distinct clades: clade A contains Cedratvirus A11, *C. zaza* and *C. lausannensis*, clade B contains *C. kamchatka*, and clade C contains Brazilian cedratvirus (Fig. S4B). This classification might be challenged as new strains will be characterized. We found that 51 of the 545 *C. kamchatka* genes were unique to this strain. According to the presence/absence of pseudogenes in the other strains, as detected using tblastn, we designed a putative evolutionary scenario for each of these genes (see Table S2). As previously discussed for pandoraviruses (37, 47), the process of *de novo* gene creation seems to participate to the shaping of Cedratviruses genomes. Interestingly, among the 51 genes unique to *C. kamchatka* only one has a clear predicted function: the ck412 DNA MTase.

### DNA methylation is widespread but unevenly distributed among giant viruses genomes

The SMRT sequencing of *C. kamchatka* and the other methylome datasets show that the various giant virus families exhibit distinct methylation features. Whereas the genomes of pithoviruses and *Mimiviridae* members are devoid of DNA modifications, those of molliviruses, pandoraviruses and cedratviruses clearly contain methylated nucleotides (Fig 1).

More specifically, *C. kamchatka* DNA is methylated at CTCG**A**G motifs (Fig. 1). Although the ck412 DNA MTase has a slightly different predicted target (CTSAG), it is probably responsible for the CTCG**A**G methylation as CTSAG motifs are not methylated (Fig. 1). *C. kamchatka* ck366 gene encodes an additional predicted FkbM domain-containing MTase for which the predicted specific target, if any, is unknown. Importantly, we found no REase associated to the *C. kamchatka* predicted MTases.

*Pithovirus sibericum* encodes two DNA MTases: pv_264 predicted to target the CTSAG motif and pv_113, an FkbM domain-containing MTase targeting an unknown site. Yet, *P. sibericum* DNA exhibit no methylated sites (including CTSAG and CTCGAG) (Fig. 1). Surprisingly, the RNA-seq data from (26) shows that both transcripts are significantly expressed all along the replication cycle (see Table S3A-B). Finally, none of the genes surrounding the MTases are predicted to encode a functional REase. Thus, *P. sibericum* MTase-like proteins do not methylate the viral DNA even though they are expressed.

Although *Megavirus vitis* encodes a type 11 domain MTase (mvi_121) and an FkbM domain-containing MTase (mvi_667) shared with Moumouvirus australiensis (ma_628), we cannot infer their DNA target specificities from sequence homology or experimental evidences since none of the two genomes appear to be methylated (Fig. 1). In addition to these MTase-like candidates, *M. vitis* and *M. australiensis* encode a 6-O-methylguanine-DNA methyltransferase (mvi_228/ma_196) probably involved in DNA repair. We also surveyed the DNA methylation of the Mimiviridae’s mobilome, namely the *Zamilon vitis* virophage, the *M. vitis* transpoviron and the *M. australiensis* transpoviron. These genomes that do not encode a DNA MTase are not methylated (Fig. 1). Collectively this data show that the DNA MTases encoded by the *Mimiviridae* infecting Acanthamoeba do not methylate the viral or mobilome DNA.

In contrast, all the surveyed Pandoraviruses’ genomes are methylated (Fig. 1). Unexpectedly they exhibit N^4^-methyl-cytosines instead of the N^6^-methyl-adenines found in *Marseilleviridae* members and Cedratviruses. The number of distinct methylated motifs is also quite variable: a single one in *P. neocaledonia*, but two in *P. celtis* and *P. quercus*, three in *P. salinus*, and up to four in *P. dulcis*. We successfully assigned each of all methylated motifs to their cognate encoded MTases. However, none appeared to be associated to a REase. In addition, we found a MTase type 25 domain-containing gene in *P. neocaledonia* (pneo_cds_672) as well as a MTase type 11 domain gene (pneo_cds_674) with orthologs in *P. dulcis*, *P. quercus* and *P. celtis* (pdul_cds_799, pqer_cds_892 and pclt_cds_906). None of them had predicted DNA targets.

Finally, the Mollivirus sibericum genome is prone to both types of modification: N^4^-methyl-cytosines and N^6^-methyl-adenines. It is methylated at the AGCA**C**T sites by the ml_135 encoded MTase and at the CTCG**A**G sites by the ml_216 MTase (Fig. 1). A third MTase encoded by this genome (ml_498) is predicted to recognize the RGATCY sites but our methylome data clearly show that they are not methylated (Fig. 1). We first suspected that the ml_498 gene was not transcribed but our transcriptomic data from (48) clearly show that ml_498 is expressed, mostly in the early phase of the infection (see Table S3C). However, the analysis of the gene structure shows a long 5’UTR suggesting a N-terminal truncation of the protein (Fig. S5A) compared to its homologs (Fig. S5B). As a single frameshift is sufficient to restore the N-terminal part of the protein, ml_498 probably underwent a recent pseudogenization.

The activity of ml_216 was further confirmed by restriction experiments showing that Mollivirus sibericum DNA is cleaved at CTCGAG sites in wild-type conditions but not after WGA (Fig. S6A). By contrast, the RGATCY sites are not protected, as expected from the ml_498 loss of function. Interestingly while the CTCGAG (ml_216), AGTACT (ml_135) and RGATCY (ml_498) sites are unmethylated in host DNA (Fig. S6). By contrast, the CCCGGG motifs are unprotected against degradation. This probably correspond to CpG methylation of the host DNA.

None of the Mollivirus sibericum MTases appear to be associated with a corresponding REase. This lack of nuclease activity was confirmed by the absence of host DNA degradation during the infection cycle (Fig. S7). One might expect that the host DNA is protected against putative CTCGAG and AGTACT targeting viral REases by endogenously encoded MTases. We exclude this possibility since host DNA is sensitive to degradation at those sites in uninfected conditions (Fig. S6B).

### The complex evolutionary history of giant viruses’ MTases

In the *Marseilleviridae* family, we observed methylation patterns typical of prokaryotic MTases. This raised the question of their evolutionary histories. In Figure S8 we now present a global phylogenetic analysis of all giant viruses’ MTases analyzed in this work (Fig. 1). They clearly do not share a common origin, and appear partitioned in 5 main groups, interspersed among bacterial and occasional amoebal homologs.

First, one observes that most viral MTases are either embedded within clusters of prokaryotic sequences (green, and red groups in Fig. S8) or constitute sister groups of prokaryotes (orange, purple and blue groups). Thus, as for Marseilleviridae these MTases are most likely of bacterial origin.

Of special interest, the tree also exhibit MTases encoded by different Acanthamoeba species (see GenBank accessions in Fig. S8), the main known hosts of giant viruses. For instance, the closest ml_135 Mollivirus MTase homologue is found in A. polyphaga, suggesting a recent exchange between virus and host. The direction of the gene transfer cannot be determined from these data. However, in the red and orange groups (Fig. S8) other acanthameoba homologues appear well nested within Pandoraviruses MTases. This supports transfers occurring from the giant viruses to the host genome. We also noticed divergent MTases attributed to various Acanthamoeba species in the purple group. However, a closer inspection of the taxonomic assignment of the corresponding contigs indicates that they are bacterial sequences, probably from the *Bradyrhizobiaceae* family (see Fig. S9). These bacteria are amoeba resistant intracellular microorganisms (56) that probably contaminated the eukaryotic host sequencing project.

The purple group also contains three orthologous MTases targeting CTCGAG sites: ml_216 from *M. sibericum*, pv_264 from *P. sibericum* and ck412 from *C. kamchatka*. These viruses belong to two distinct viral families but infect similar hosts. It is thus likely that these MTases were recently exchanged between these viruses. In the green group there are also three orthologous MTases from distinct viral families: pdul_cds_639 and ppam_cds_578 from *P. dulcis* and *P. pampulha* respectively, as well as ml_498 from *M. sibericum*. The two pandoraviruses are closely related with an average of 83% sequence identity between shared proteins. As this MTase is not found in other pandoraviruses (Fig. S10), a gene exchange might have occurred between the *P. pampulha*-*P. dulcis* ancestor and *M. sibericum*.

### Selection pressure acting on giant viruses’ MTases

Following the above phylogenetic analysis of the giant viruses MTases, we investigated the selection pressure acting on them. We first noticed that some MTases were conserved for long periods of time in various viral families. The **C**TCGAG and C**C**CGGG targeting MTases, most likely gained by a Pandoravirus ancestor, remain present in most of the extant Pandoraviruses (Fig. S10). Likewise, the Marseilleviridae R-M MTase and the C**C**TNAGG Pandoravirus targeting MTase were kept in almost all members of their respective clades (Fig. 2 and Fig. S10). By contrast, the ml_498 *M. sibericum* MTase was found to be recently pseudogenized (Fig. S5). In addition, the pino_cds_419 gene from *P. inopinatum* and the ppam_cds_578 from *P. pampulha* are most likely truncated pseudogenes, even though we do not have SMRT data to confirm their loss of function.

We then computed the ratios (ω) of non-synonymous (dN) to synonymous (dS) substitution rates to quantify the selection pressure acting on the MTases. The ω of MTases with predicted targets were calculated using Codeml (57) according to three different models (see Methods). We then selected the best fitted models using likelihood ratio tests (LTR) to determine whether ω were significantly different from one. As shown in Table S4, the majority (15/24) of the MTases had a ω significantly smaller than one and the rest could not be statistically distinguished from neutral evolution. This indicates that most giant viruses MTases are under purifying selection.

## Discussion

Following the initial description of Mimivirus (21), the last decade has seen an acceleration in the pace of discovery of giant viruses, now distributed in multiple different families (20, 23, 26, 27, 48, 58–60), both thanks to the physical isolation of new specimens and to the rapid accumulation of metagenomics data (31, 34, 36). Although the number of genomic sequences steadily increased during this period, the epigenomic status of giant viruses remained virtually unknown. Yet, the presence of numerous predicted DNA modification functions in their gene contents, as well as histone homologs in some of them (61, 62), suggest that epigenetic may have a general impact on giant viruses’s fitness, most likely through virus-virus and host-virus interactions. Here, we presented the first investigation of the DNA methylome of a large diversity of giant viruses using SMRT sequencing. Our analyses reveal that DNA methylation is widespread as it was detected in four of the 5 distinct giant viruses’ families tested (Mimiviridae, Cedratviruses, Molliviruses, Pandoraviruses and Marseilleviridae). The recent advances in SMRT sequencing of metagenomes (44) will probably soon enable the survey of cultivation-independent giant viruses (31, 63) and comfort this observation.

Our detailed investigation of the DNA MTase gene contents first confirmed the ubiquity of DNA methylation in giant viruses. We identified homologs of these enzymes in all analyzed viruses, with the exception of the Mimiviridae’s mobilome. Although widely present in giant viruses, the number of encoded MTases (and targeted sites) is unevenly distributed. It ranges from a single one in Melbournevirus and *M. australiensis*, up to five in *P. dulcis*. Even within the same family, such as the the Pandoraviruses, the number of encoded DNA MTases is variable. Furthermore, the number of occurrences of each methylated sites per genome is highly variable, from 4 (C**C**TNAGG in *P. quercus*) to 850 (CTCG**A**G in *M. sibericum*) (Fig. 1). The range of relative frequency of these sites is even larger from 10^−6^ for C**C**TNAGG in *P. quercus* up to 10^−3^ for the G**A**TC motif in Melbournevirus. Therefore, as already noticed in prokaryotes (42–44), DNA methylation is widespread but has a patchy distribution in giant viruses.

The non-uniformity of DNA methylation in giant viruses is partially explained by the loss-of-function of some encoded MTases. For instance, the *M. sibericum* ml_498 MTase lacks a conserved (D/N/S)PP(Y/F) motif, involved in the formation of a hydrophobic pocket that binds the targeted nucleotide (64). This was probably caused by a recent frameshift mutation in the 5′ region of the gene. Another case is the *P. sibericum* pv_264 MTase also unable to methylate viral DNA, although the protein does not seem to be truncated. Here, uncharacterized mutations in a critical part of the protein might be at play.

As expected from their enzymatic specificities, giant viruses’ MTases are all of bacterial origins (Fig. S8). More surprisingly, we found that some of them were transferred from giant viruses (mostly Pandoraviruses) to Acanthamoeba genomes. It remains to be determined whether this is an evolutionary dead end or if the transferred enzymes are still active in the host. Although host-to-virus gene exchanges are traditionally deemed more frequent than virus-to-host transfers (65, 66), we previously noticed that this might not be true in Pandoraviruses (37). The picture is even more complex concerning MTases transferred between viruses, as illustrated by the orthologous MTases identified in *C. kamchatka* and *M. sibericum*, two viruses from different families. If we previously noticed that some genes might be swapped between strains of pandoraviruses (67), the present case involves an exchange between viruses from totally different families only sharing the Acanthamoeba host. In the recently discovered *Mollivirus kamchatka* (68), a MTase without ortholog in *M. sibericum* was probably acquired from a Pandoravirus. In prokaryotes, the analysis of the co-occurrence of R-M systems and genetic fluxes between bacteria revealed that genetic exchanges are favored between genomes that share the same R-M systems, regardless of their evolutionary distance (69). A similar phenomenon might be at play between Molliviruses and Pandoraviruses, and partially explains their shared gene content (68). Previous analyses have also suggested that amoeba act as a genetic melting pot between intracellular bacteria (70, 71), a concept that should now be extended to include amoeba-infecting viruses.

Our global survey of the DNA methylome of giant viruses revealed unexpected features. Besides the N^6^-methyl-adenine modifications identified in the Melbournevirus, *C. kamchatka* and *M. sibericum,* we unveiled unexpected N^4^-methyl-cytosines in the genomes of *M. sibericum* and all tested pandoraviruses. To our knowledge, these are the first of such modifications reported for eukaryotic viruses. Some chloroviruses contain large amounts of N^6^-methyl-adenines and 5-methyl-cytosines (72, 73) also found in other eukaryotic viruses such as herpesviruses (12, 74) and members of the Iridoviridae (12, 75), and Adenoviridae (12, 76) families. However, N^4^-methyl-cytosine modifications were until now thought to be restricted to the prokaryotes and their viruses.

Another unexpected finding is the discovery of different types of modifications of the same CTCGAG site in giant viruses. If *C. Kamchatka* and *M. sibericum* exhibit CTCG**A**G motifs (with N^6^-methyl-adenines), the pandoraviruses exhibit **C**TCGAG with N^4^-methyl-cytosines (Fig. 1). The corresponding MTases belong to two distinct phylogenetic groups (green and orange groups in Fig. S8) and were acquired from distinct prokaryotes. Structural studies will be needed to elucidate how these MTases differently methylate the same DNA motif.

It was recently discovered that the AGCT tetramer is specifically eliminated from the Pandoraviruses’ genomes, in particular the ones belonging to the A-clade (77). The evolutionary mechanism causing the elimination of this motif is still mysterious. One of the rejected hypothesis was that a R-M system targeting AGCT sites could be involved (77). Our methylome data consistently confirm that the AGCT motif is not methylated in the pandoraviruses of neither clade A nor B (Fig. S11).

Our digestion experiments revealed the presence N^6^-methyl-adenine-modified G**A**TC sites in the genome of *A. castellanii* (Fig. 3). N^6^-methyl-adenines were long thought to be restricted to prokaryotes, until several studies showed that some eukaryotes are also subject to these modifications (78–80). In Chlamydomonas for instance, the N^6^-methyl-adenines preferentially localize at the vicinity of transcription start sites, in the nucleosome free regions, to mark transcriptionally active genes (79). Since the *A. castellanii* genome contains a mixture of methylated and unmethylated GATC sites we expect that similar biased modification patterns could be revealed by SMRT sequencing.

Most of the MTases (16 over 18) which had a testable (i.e. predicted) target site were found to be functional (Fig. 1). This either suggests that they were recently acquired, or that they were conserved because they increased the recipient viruses’ fitness. Several evidences favor the second hypothesis. First, we found several of them conserved in entire clades (Fig. 2, Fig. S10), indicating that they were retained throughout the family’s radiation. Secondly, most of them are under purifying selection (Table S4).

We observed that the complete R-M systems found in the *Marseilleviridae* members were phylogenetically related to that of the Chloroviruses, in which it functions as a host DNA recycling mechanism (81–83). There was thus a possibility that the Marseilleviridae R-M system could play a similar role. However, our data clearly refute this hypothesis: even though we found that the host DNA remains vulnerable at the GATC site targeted by the Marseilleviridae REase (Fig. 3), the actual infection did not induce its degradation (Fig. S3). This result is coherent with what we know about the replication cycle of these viruses (53, 61, 62). Even if Melbournevirus temporarily requires nucleus functions to initiate its replication cycle (53), most of it then proceeds in the cytoplasm, without contact with the host DNA (53). This supports the non-involvement of Marseilleviridae R-M systems, as well as other encoded REases, in the recycling of the host DNA.

By contrast, our results revealed that REases corresponding to the Marseilleviridae R-M system could degrade the DNA of other marseilleviruses devoid of the same system. As previously proposed for Chloroviruses (19, 84) this suggests that the Marseilleviridae R-M systems are involved in the exclusion of other viruses in cases of multiple infections. Indeed, Acanthamoeba can be infected by a wide variety of giant viruses (20, 85). In this context, a R-M system becomes an efficient way for a virus to fight against competitors and increase its fitness. In addition, Acanthamoeba feed on bacteria and are the reservoir of many intracellular bacteria, some of them remain as cytoplasmic endosymbionts (56, 86). The Marseilleviridae R-M systems could thus be involved in recycling the DNA of these intracellular parasites.

In line with such putative role in pathogen exclusion, we found a congruent pattern of expression of the Melbournevirus R-M system REase and MTase. The REase is first transcribed in the early phase of the infection cycle, when it could degrade the DNA of eventual co-infecting bacteria or viruses. The MTase is then transcribed 15min later and the enzyme finally packaged in the virion, where it could protect the viral DNA from the REase, pending the next infection.

Besides Marseilleviridae family members, giant viruses for which we identified MTases and observed DNA methylation do not seem to encode cognate REases. Such so-called orphan MTases are common in bacteria where they regulate various biological processes, such as replication initiation, mismatch repair or gene expression (7–11). Accordingly, the targeted genomic positions are not uniformly distributed, with hotspots and coldspots of fully-, hemi- and unmethylated sites. In contrast, the DNA methylation of giant viruses’ genomes exhibits unimodal distributions of IPDr values, and globally uniform positioning of the corresponding motifs (Fig. S12). In the case of bacterial orphan MTases involved in gene regulations the methylated sites tend to be located in the upstream non-coding regions of the regulated genes (42). This is not true for giant viruses where methylated motifs are not enriched in intergenic regions (p-values > 0.05, see Methods).

How could we then interpret the presence of orphan MTases in giant viruses? First, their role could be to protect the viral genome from its digestion by cognate REases from other Acanthamoeba-co-infecting bacteria or viruses harboring complete R-M systems. This would explain the tendency for some viruses, such as *P. dulcis*, to accumulate functional MTases in their genomes. In absence of the corresponding REases in the environment, the selection pressure would be relaxed on less solicited MTases leading to their pseudogenization (Table S4).

Giant viruses of the Mimiviridae family are involved in complex network of interactions with the cellular host, the virophages, and the transpovirons (45, 87–90). It has been proposed that R-M systems could be used as an anti-virophage agent in these multipartite systems (91). Accordingly, we investigated the role of DNA methylation in these cross talks. Our analysis of Acanthamoeba-infecting Mimiviridae members do not currently support this view, as DNA methylation do not seem to be a key player in this network (Fig 1). However, systems involving other hosts, such as the Cafeteria roenbergensis-CroV-Mavirus trio (25, 89, 92), might depend on DNA methylation to regulate their intricate interactions. DNA methylation is also a key factor in the switch between latent and integrated forms of some viruses (12). In the Cafeteria roenbergensis-CroV-Mavirus context, one might wonder about the role DNA methylation could play in the maintenance of the host integrated mavirus provirophage, or in its awakening in the presence of CroV infections.

R-M systems provide the carrying bacteria an immediate protection against the most lethal bacteriophages present it its environment (6). Our work suggests that DNA methylation is equally important in giant viruses and involved in several types of interactions depending on the presence or absence of REase activity, and on the strictly-cytoplasmic or nucleus dependency of their replication cycle. In Chloroviruses, R-M systems offer a way to attack host DNA and exploit its nucleotide pool (83). Our work on Marseilleviridae now suggest that they act as an offensive weapon against competing pathogen. By contrast, the many orphan MTases found in giant viruses are potential self-defense weapons against other pathogens bearing active R-M systems with similar targets. Therefore, DNA methylation might allow giant viruses to face the fierce battles taking place in their amoebal hosts. In that context, it seems odd that the most frequent giant viruses in the environment that we studied, the *Mimiviridae*, are apparently devoid of DNA methylation. But one might remember that bacteria developed different lethal weapons to survive phage invasion: R-M and CRISPR systems. The many other REases found in the giant viruses’ genome could thus be involved in other competition/resistance processes such as the “Mimivire” system suggested to be directed against the parasitic virophage competing with some *Mimiviridae* members for its replication in Acanthamoeba (93–95). Alongside toxin-antitoxin systems (91, 96), Mimivire or other yet to be discovered CRISPR-like systems, DNA methylation might be part of the giant viruses’ arsenal to cope with their numerus competitors.

## Methods

### Cedratvirus kamchatka characterization

#### Isolation and purification

*Cedratvirus kamchatka* was isolated from muddy grit collected near a lake at kizimen volcano, Kamchatka (Russian Federation N 55° 05′ 50 E 160° 20′ 58). The sample was resuspended in Phosphate Buffer Saline containing Ampicillin (100µg/ml), Chloramphrenicol (30µg/ml) and Kanamycin (25µg/ml) and an aliquot was incubated with *Acanthamoeba castellanii Neff* (ATCC30010TM) cells (2 000 cells per cm^2^) adapted to 2.5µg/mL of Amphotericin B (Fungizone), in protease-peptone-yeast-extract-glucose (PPYG) medium. Cultures exhibiting infectious phenotypes were recovered, centrifuged 5 min at 500g to eliminate the cells debris and the supernatant was centrifuged for 1 hour at 16,000 g at room temperature. T75 flasks were seeded with 60,000 cells/cm^2^ and infected with the resuspended viral pellet. After a succession of passages, viral particles produced in sufficient quantity were recovered and purified.

The viral pellet was resuspended in PBS and loaded on a 1.2 to 1.5 density cesium gradient. After 16 h of centrifugation at 200,000 g, the viral disk was washed 3 times in PBS and stored at 4°C.

#### Genome sequencing and assembly

The genomic DNA was recovered from 2×10^10^ purified particles resuspended in 300µL of water incubated with 500µL of a buffer containing 100 mM Tris-HCl pH 8, 1.4 M NaCl, 20 mM Na2EDTA, 2% (w/v) CTAB (Cetyltrimethylammonium bromide), 6mM DTT and 1 mg/mL proteinase K at 65°C for 90 min. After treatment with 0.5 mg/mL RNase A for 10min, 500µL of chloroform was added and the sample was centrifuged at 16,000 g for 10 min at 4°C. One volume of chloroform was added to the supernatant and centrifuged at 16,000 g for 5 min at 4°C. The aqueous phase was incubated with 2 volumes of precipitation buffer (5 g/L CTAB, 40 mM NaCl, pH 8) for 1 h at room temperature and centrifuged for 5 min at 16,000 g. The pellet was then resuspended in 350µL of 1.2M NaCl and 350 µL of chloroform was added and centrifuged at 16,000 g for 10 min at 4°C. The aqueous phase was mixed to 0.6 volume of isopropanol, centrifuged for 10 min at room temperature and the pellet was washed with 500 µL 70% ethanol, centrifuged again and resuspended with nuclease-free water.

C. kamchatka DNA was sequenced using PacBio SMRT technology, resulting in 868 Mb of sequence data (76,825 reads). SMRT reads were filtered using the SMRTanalysis package version 2.3.0. We then used Flye (97) version 2.4.2 to perform the *de novo* genome assembly with the “pacbio_raw” and “g=450000” parameters. The assembly resulted in two distinct contigs (one of 466,767 nt and a second of 6,049 nt). A pulsed field gel electrophoresis (PFGE) confirmed the genome size and its circular structure (Fig. S4A). The smaller contig, not seen on the PFGE, potentially corresponds to an assembly artefact. Finally the assembly was subsequently polished using the Quiver tool from SMRTanalysis.

#### Genome annotation

The C. kamchatka gene annotation was performed using GeneMarkS (98) with the “—virus” option. Only genes predicted to encode proteins of at least 50 amino acids were kept for functional annotation. These proteins were aligned against the NR and Swissprot databases using BlastP (99) (with an E-value cut-off of 10^−5^) and submitted to CD search (49), to InterProScan (100) with the Pfam, PANTHER, TIGRFAM, SMART, ProDom, ProSiteProfiles, ProSitePatterns and Hamapts databases, and to Phobius (101). The genome was then manually curated according to these data.

### Pithovirus sibericum and Mollivirus sibericum resequencing

P. sibericum and M. sibericum genomic DNAs were extracted from 2×10^10^ purified particles using the the PureLink Genomic DNA Extraction Mini Kit (Thermo Scientific) according to the manufacturer protocol. For P. sibericum, we performed 2 successive purifications, and added 10 mM DTT in the lysis buffer for the first one.

The sequencing of P. sibericum and M. sibericum performed using PacBio SMRT technology resulted in respectively 779 Mb (143,675 reads) and 371 Mb (66,453 reads) of sequence data. A second SMRT sequencing was performed on P. sibericum and M. sibericum viral DNA after WGA amplification using the Illustra GenomiPhi V2 DNA Amplification kit (GE Healthcare) according to the manufacturer instructions. This resulted in 632 Mb (78,685 reads) and 346 Mb (55,797 reads) of sequences for P. sibericum and M. sibericum WGA amplified DNA.

### Identification of methylated motifs

Methylated motifs were identified using the “Modification and motif analysis” module from the SMRTanalysis package using the datasets described in Table S1 and the corresponding reference genome sequences. In addition we used the per-base resolution file of IPD ratios calculated by SMRTanalysis to compute the global IPDr of detected motifs and predicted MTases targets.

### Annotation of MTases, REases and target prediction

We analyzed the MTases and REases encoded in giant viruses genomes based on published annotations when available, as well as a combination of CD search and Interproscan protein domain search tools analyses (49, 50). In addition, we performed blast alignments against the REbase database (51) to support the annotations and to predict their targets when possible.

### Phylogenies

All the phylogenies were calculated from the protein multiple alignments of homologous sequences computed using Expresso (102), Mafft (103), Mcoffee (104) and Clustal Omega (105). We next selected the best multiple alignment using TrimAI (106). For phylogenies based on single genes we used IQtree (107) to compute the best model and the tree. Bootstrap values were computed using the UFBoot method (108). Phylogenies of viral families were based on the multiple alignments of strictly conserved single copy genes. These groups of orthologs were found using the OrthoFinder algorithm (109). Next we used IQtree (107) to compute the tree using the best model of each partitioned alignment.

### Selection pressure measurements

We performed codon-based multiple alignments of each subgroup of giant viruses MTases (Fig. S8) using protein multiple alignments (see Phylogenies Methods) and nucleotide sequences. The ω were compute for each gene of interest using Codeml (57) through the ETE framework (110). Three models were used: the “M0” model which considers a unique ω for the whole tree, the “B_free” model which assigns two distinct ω (one for the branch of interest and one for the rest of the tree), and the “B_neut” model similar to “B_free” but with a fixed ω=1 for the gene of interest. LTR tests were then performed to decide whether ω were significantly different from one.

### DNA restriction experiments

Mollivirus sibericum, Melbournevirus and Noumeavirus genomic DNA were extracted using the PureLink Genomic DNA Extraction Mini Kit (Thermo Scientific) according to the manufacturer protocol. A. castellanii genomic DNA was extracted using the Wizard^®^ Genomic DNA Purification Kit (Promega). The DNA were digested with 10 units of the appropriate restriction enzymes (New England Biolabs) for 1 h at 37°C and loaded on a 1% agarose gel.

### Mollivirus sibericum-infected A. castellanii cells DNA extraction

A. castellanii overexpressing the ml_216-GFP fusion and wild-type cells were grown in T25 flasks. They were infected with M. sibericum at MOI 100. After 1 h of infection at 32°C, cells were washed 3 times with PPYG to eliminate the excess of viruses. For each infection time (1h to 6h), a T25 flask was recovered and cells were centrifuged for 5min at 1,000 g. DNA was extracted using the Wizard^®^ Genomic DNA Purification Kit (Promega) according to the manufacturer’s protocol and loaded on a 0.8% agarose gel.

### Melbournevirus R-M system MTase and REase expression timing

A. castellanii cells were grown in T25 flasks and infected with Melbournevirus at MOI 50. After 15 min of infection at 32°C, cells were washed 3 times with PPYG to eliminate the excess of viruses. For each infection time (15min to 5h), a T25 flask was recovered and cells were centrifuged for 5min at 1,000 g. RNA was extracted using the RNeasy Mini kit (QIAGEN) according to the manufacturer’s protocol. Briefly, cells were resuspended in the provided buffer and disrupted by −80 °C freezing and thawing, and shaken vigorously. Total RNA was eluted with 50 μl of RNase free water. Total RNA was quantified on the nanodrop spectrophotometer (Thermo Scientific). Poly(A) enrichment was performed (Life Technologies, Dynabeads oligodT_25_) and first-strand cDNA poly(A) synthesis was performed with the SmartScribe Reverse Transcriptase (Clontech Laboratories) using an oligo(dT)_24_ primer and then treated with RNase H (New England Biolabs). For each time point, PCR reactions were performed using MEL_015 and MEL_016 specific primers and one unit of Phusion DNA Polymerase (Thermo Scientific) in a 50 μl final volume.

### Motif enrichment in intergenic regions

Analysis of motif enrichment in intergenic regions was done by computing the number of motifs identified in these regions and in coding regions. We next shuffled coding coordinates 1,000 times and calculated the same values. Z-scores were calculated from these randomizations and transformed in empirical p-values. We excluded the CCTNAGG motifs from this calculation as there were not enough occurrences of this motif (Fig. 1) to compute accurate p-values.

Cedratvirus kamchatka genome GenBank accession: XXXXX.

---

**Figure S1. Control experiment based on SMRT sequencing of Mollivirus sibericum and Pithovirus sibericum native and WGA amplified DNA**

The encoded DNA MTases of both viruses are shown along with their predicted target motifs. Modified nucleotides are indicated in bold. The median IPDr profiles of the MTases targets (in gray) and their surrounding 20 nucleotides on each side are shown. Red bars correspond to positions with significantly high IPDr values. Error bars correspond to 95% intervals based on bootstrapping. The median IPDr values of native and WGA amplified DNA are shown side by side.

**Figure S2. Phylogenies of the Marseilleviridae R-M system MTase and REase**

**A**) Phylogenetic tree of the mel_016 Melbournevirus MTase along with viral and prokaryotic homologs. The tree was calculated using the LG+F+I+G4 model based on a protein sequence alignment of 321 informative sites. Bootstrap values were computed using the UFBoot (108) method from IQtree (107). The GenBank accessions and taxonomic assignations extracted from GenBank entries are shown. Virion symbols highlight viral sequences that are classified in their corresponding family using a color code described in the inset. Prokaryotic sequences are not color coded. It is worth noting that the Iridovirus sequence comes from a metagenomics assembly (33) and that its taxonomic assignment is uncertain when considering the best blast matchs againt NR of the ORFs encoded in the corresponding genomic contig (Fig. S9A). Likewise, the Rhodobacteraceae bacterial PQM57433.1 sequence also recovered from metagenome assemblies (111) is probably of viral origin (Fig. S9B). **B**) Similar phylogenetic tree of the mel_015 Melbournevirus MTase. The tree was calculated using the VT+G4 model based on a protein sequence alignment of 188 informative positions.

**Figure S3. kinetics of host DNA degradation upon Melbournevirus and Noumeavirus infections**

Host DNA is not degraded during Melbournevirus and Noumeavirus infections. Times (post infection) are listed on the top of the figure. NI corresponds to non-infected.

**Figure S4. Characterization of the Cedratviruse kamchatka genome**

**A**) Pulse Field Gel Electrophoresis of the C. kamchatka genome. The first and last lanes correspond to DNA ladders. Undigested and NotI digested DNA are shown. As expected from its circular structure the undigested DNA migrates higher than the assembled genomic sequence length. The NotI digested DNA, predicted to be cleaved at a single position migrates at the expected position. **B**) Phylogenetic tree of the completely sequenced genomes of Cedratviruses and Pithoviruses based on the protein sequences alignments of 134 strictly conserved single copy orthologues. The tree was calculated using the best model of each partitioned alignment as determined by IQtree (107). Bootstrap values (n=5,000) were computed using the UFBoot (108) method from IQtree (107). The average amino acid identity matrix was computed using CompareM (112) on the shared ORFs.

**Figure S5. Gene structure of the Mollivirus sibericum ml_498 gene**

**A)** The structure of the ml_498 gene is shown. Protein coding sequence is in blue and UnTranslated Regions (UTRs) in gray. The transcriptomic RNA-seq data from (48) was mapped to the genome and read density is shown in gray at early, intermediate and late infection time points (see 48 for details). **B)** Protein sequence multiple alignment of ml_498 and homologues (named according to their GenBank accession) produced using the Expresso tool (102). The multiple alignment includes the ml_498 protein sequence as well as a version of the protein sequence were a frameshift in the 5’ region of the gene was corrected (ml_498_frameshited). Sequence conservation is highlighted in blue. The red box encompasses a highly conserved xPPY functional motif present in DNA MTases.

**Figure S6. Host and Mollivirus sibericum DNA methylation assessed by restrictions**

A) Agarose gel electrophoresis analysis of M. sibericum and host DNA after digestion with restriction enzymes targeting the same sites as the encoded M. sibericum MTases. Restriction experiments were performed on A. castellanii DNA, M. sibericum DNA as well as M. sibericum DNA following whole genome amplification. The theoretical number of fragments expected from the number of occurrences of the motifs in the genomic sequences are noted at the bottom of the figure. B) A. castellanii host DNA restriction with REases sharing the same targets than ml_216 and ml_135 in wild type conditions and after expression of a GFP protein or a GFP fused with ml_216.

**Figure S7. Kinetics of host DNA degradation upon Mollivirus sibericum infections**

Host DNA was not degraded during M. sibericum infection. Times (post infection) are listed on the top of the figure. NI corresponds to non-infected.

**Figure S8. Phylogenetic tree of the giant viruses MTases**

Phylogenetic tree of the giant viruses’ MTases along with prokaryotic and eukaryotic homologues. Viral sequences are highlighted using a virion symbol, eukaryotic sequences are highlighted in yellow and prokaryotic sequences are not color coded. The tree was computed using the LG+R6 model from a multiple alignment of 678 informative sites. Bootstrap values were computed using the UFBoot (108) method from IQtree (107) and reported to the branches according to the color code described in the inset. The GenBank accessions and taxonomic assignations extracted from GenBank entries are shown. The tree is divided into 5 color coded subgroups.

**Figure S9. Taxonomic assignment of metagenomics and Acanthamoeba contigs**

For each contig (A-G) the taxonomic assignment of the best blastp matchs (with E-value < 10^−5^) against the NR database of all the encoded ORFs is summarized in a Krona chart (113). The GenBank accessions of the contigs are displayed as well as the number of ORFs with a significant match and the number of ORFs without match (ORFans).

**Figure S10. Presence/absence of Pandoraviruses MTases mapped onto the Pandoraviruses phylogeny**

The phylogenetic tree was computed using the protein sequence alignments of 375 strictly conserved single copy orthologues. The tree was calculated using the best model of each partitioned alignment as determined by IQtree (107). Bootstrap values were computed using the UFBoot (108) method from IQtree (107) but not reported as they were all equal to 100%. For each sub-group of Pandoraviruses encoded MTases the presence/absence of the gene is shown with its predicted sequence target using the same colors as in Fig. S8. A filled circle means that a MTase of a given sub group is encoded, while an empty circle means that it is absent. Bold circles highlight MTases for which SMRT data is available while shaded circles depict MTases with no available SMRT data. Gray circles show potential pseudogenes. The colored arrows point to the most parsimonious timings of the MTase acquisition for each sub-group.

**Figure S11. Methylation status of the underrepresented AGCT Pandoraviruses’s tetramer**

The median IPDr profiles of the AGCT tetramer (highlighted in gray) and the surrounding 20 nucleotides on each side are shown. Error bars correspond to 95% intervals based on bootstrapping. P. salinus and P. celtis belong to clade A and P. neocaledonia to clade B (67).

**Figure S12. Distributions of IPDr values and motifs along the genomes**

For each virus and methylated motif pair, we calculated the IPDr values distributions (left graph). We also computed the empirical cumulative distribution of the number of motif occurrence along the genome (right graph). The Kolmogorov-Smirnov statistical test was used to determine if the motifs were uniformly distributed along the genome. The CCTNAGG motif was excluded from these analyses as there are not sufficient occurrences of this motif in the genomes (see Fig. 1).

**Table S1. Datasets references**

**Table S2. Protein-coding genes unique to Cedratvirus kamchatka**

**Table S3. RNA-seq transcriptomic data of Pithovirus sibericum and Mollivirus sibericum MTases**

**Table S4. dN/dS ratios of MTases**

## Acknowledgments

We are deeply indebted to our volunteer collaborator Alexander Morawitz for collecting the Kamchatka soil samples. We thank the PACA Bioinfo platform for computing support. This project has received funding from an innovation program (grant agreement No 832601), from the FRM prize “Lucien Tartois”, and from CNRS (PRC1484-2018) to C. Abergel. The funding bodies had no role in the design of the study, analysis, and interpretation of data and in writing the manuscript.

