## Supplementary material for "The DNA Methylation Landscape of Giant Viruses": TableS1

### Table S1.

#### A. Datasets

| Virus | Reference genome Genbank ID | Reference of the PacBio SMRT data |
|---|---|---|
| Megavirus vitis | MG807319.1 | Jeudy et al. doi: 10.1038/s41396-019-0565-y |
| Moumouvirus australiensis | MG807320.1 | |
| Zamilon zitis | MG807318.2 | |
| Megavirus vitis transpoviron | MG807316.1 | |
| Moumouvirus australiensis transpoviron | MG807317.1 | |
| Melbournevirus | KM275475.1 | Philippe N et al. doi: 10.1126/science.1239181 |
| Pandoravirus dulcis | KC977570.1 | |
| Pandoravirus salinus | KC977571.1 | |
| Pandoravirus quercus | MG011689.1 | Legendre M et al. doi: 10.1038/s41467-018-04698-4 |
| Pandoravirus neocaledonia | MG011690.1 | |
| Pandoravirus celtis | MK174290.1 | Legendre M et al. doi: 10.3389/fmicb.2019.00430 |
| Mollivirus sibericum | KR921745.1 | This study |
| Pithovirus sibericum | KF740664.1 | This study |
| Cedratvirus kamtchatka | This study | This study |

#### B. *Marseilleviridae* members complete genomes

| **Virus** | **Reference genome Genbank ID** |
|---|---|
| Melbournevirus | KM275475.1 |
| Cannes 8 virus | KF261120.1 |
| Marseillevirus | GU071086.1 |
| Marseillevirus shanghai 1 | MG827395.1 |
| Tokyovirus A1 | AP017398.1 |
| Kurlavirus | KY073338.1 |
| Noumeavirus | KX066233.1 |
| Port-Miou virus | KT428292.1 |
| Lausannevirus | HQ113105.1 |
| Brazilian marseillevirus | KT752522.1 |
| Insectomimevirus | KF527888.1 |
| Tunisvirus | KF483846.1 |
| Golden marseillevirus | KT835053.1 |
