## Supplementary material for "The DNA Methylation Landscape of Giant Viruses": TableS2

### Table S2. Protein-coding genes unique to Cedratvirus kamchatka

| Gene | Predicted function | Putative evolutionary scenario |
|---|---|---|
| ck112 | Hypothetical protein | *de novo* creation (or HGT from an unknown organism) and loss in clade A |
| ck113 | Hypothetical protein | Uncertain |
| ck114 | Hypothetical protein | *de novo* creation or HGT from an unknown organism |
| ck12 | Hypothetical protein | Putative HGT |
| ck137 | Hypothetical protein | *de novo* creation or HGT from an unknown organism |
| ck14 | Hypothetical protein | *de novo* creation (or HGT from an unknown organism) and loss in clade A and C |
| ck153 | Hypothetical membrane protein | *de novo* creation or HGT from an unknown organism |
| ck154 | Hypothetical protein | *de novo* creation (or HGT from an unknown organism) and loss in clade A and C |
| ck155 | Hypothetical protein | Putative HGT |
| ck16 | Hypothetical protein | *de novo* creation or HGT from an unknown organism |
| ck17 | Hypothetical protein | *de novo* creation or HGT from an unknown organism |
| ck170 | Hypothetical protein | *de novo* creation (or HGT from an unknown organism) and loss in clade A |
| ck18 | Hypothetical membrane protein | *de novo* creation or HGT from an unknown organism |
| ck207 | Hypothetical protein | *de novo* creation (or HGT from an unknown organism) and loss in clade A |
| ck208 | Hypothetical membrane protein | *de novo* creation (or HGT from an unknown organism) and loss in clade A |
| ck209 | Hypothetical membrane protein | *de novo* creation (or HGT from an unknown organism) and loss in clade A |
| ck210 | Hypothetical membrane protein | *de novo* creation or HGT from an unknown organism |
| ck211 | Hypothetical protein | *de novo* creation (or HGT from an unknown organism) and loss in clade A |
| ck213 | Hypothetical protein | *de novo* creation or HGT from an unknown organism |
| ck226 | Hypothetical protein | *de novo* creation or HGT from an unknown organism |
| ck236 | Hypothetical membrane protein | Putative HGT |
| ck237 | Hypothetical membrane protein | *de novo* creation or HGT from an unknown organism |
| ck238 | Hypothetical protein | *de novo* creation (or HGT from an unknown organism) and loss in clade A |
| ck266 | Hypothetical protein | *de novo* creation or HGT from an unknown organism |
| ck278 | Hypothetical protein | *de novo* creation or HGT from an unknown organism |
| ck312 | Hypothetical protein | *de novo* creation (or HGT from an unknown organism) and loss in clade A |
| ck345 | Hypothetical protein | Putative HGT |
| ck360 | Hypothetical protein | *de novo* creation (or HGT from an unknown organism) and loss in clade A |
| ck412 | DNA-Adenine methyltransferase | HGT |
| ck421 | Hypothetical protein golgin subfamily A member 6-like | Putative HGT and loss in clade A |
| ck423 | Hypothetical protein | *de novo* creation (or HGT from an unknown organism) and loss in clade A and C |
| ck469 | Hypothetical protein | *de novo* creation or HGT from an unknown organism |
| Ck471 | Hypothetical membrane protein | *de novo* creation (or HGT from an unknown organism) and loss in clade A and C |
| ck472 | Hypothetical protein | *de novo* creation (or HGT from an unknown organism) and loss in clade A |
| ck473 | Hypothetical membrane protein | *de novo* creation (or HGT from an unknown organism) and loss in clade A |
| ck474 | Hypothetical membrane protein | *de novo* creation (or HGT from an unknown organism) and loss in clade A |
| ck476 | Hypothetical protein | *de novo* creation (or HGT from an unknown organism) and loss in clade A |
| ck477 | Hypothetical membrane protein | *de novo* creation (or HGT from an unknown organism) and loss in clade A |
| ck478 | Hypothetical membrane protein | *de novo* creation (or HGT from an unknown organism) and loss in clade A and C |
| ck479 | Hypothetical protein | *de novo* creation (or HGT from an unknown organism) and loss in clade A |

| | | |
|---|---|---|
| ck480 | Hypotetical membrane protein | *de novo* creation (or HGT from an unknown organism) and loss in clade A |
| ck481 | Hypothetical protein | *de novo* creation (or HGT from an unknown organism) and loss in clade A |
| ck52 | Hypothetical protein | *de novo* creation or HGT from an unknown organism |
| ck522 | Hypothetical protein | *de novo* creation or HGT from an unknown organism |
| ck523 | Hypothetical protein | *de novo* creation or HGT from an unknown organism |
| ck524 | Hypothetical protein | *de novo* creation (or HGT from an unknown organism) and loss in clade A |
| ck525 | Hypothetical membrane protein | *de novo* creation or HGT from an unknown organism |
| ck526 | Hypothetical protein | *de novo* creation or HGT from an unknown organism |
| ck90 | Hypothetical protein | *de novo* creation or HGT from an unknown organism |
| ck95 | Hypothetical protein | *de novo* creation or HGT from an unknown organism |
| ck96 | Hypothetical membrane protein | Putative HGT |
