## Supplementary material for "The DNA Methylation Landscape of Giant Viruses": TableS3

### Table S3A. pv_264 transcript expression levels

| | pv_264 mapped reads | FPKM | Gene expression rank (among 467 P. sibericum genes) |
|---|---|---|---|
| Non infected | 0 | 0 | NA |
| 4h pi | 6 | 62.7 | 181 |
| 11h pi | 5 | 82.9 | 221 |
| 16h pi | 8 | 454.2 | 107 |

### Table S3B. pv_113 transcript expression levels

| | pv_113 mapped reads | FPKM | Gene expression rank (among 467 P. sibericum genes) |
|---|---|---|---|
| Non infected | 6 | 0 | NA |
| 4h pi | 65 | 78.2 | 41 |
| 11h pi | 38 | 72.4 | 51 |
| 16h pi | 15 | 103.2 | 67 |

### Table S3C. ml_498 transcript expression levels

| | ml_498 mapped reads | FPKM | Gene expression rank (among 523 M. sibericum genes) |
|---|---|---|---|
| 30min-1h-2h pi | 8982 | 168.7 | 149 |
| 3h-4h-5h pi | 15593 | 252.2 | 383 |
| 6h-7h-9h pi | 7187 | 113.6 | 398 |
