## Supplementary material for "The DNA Methylation Landscape of Giant Viruses": TableS4

### Table S4. dN/dS ratios of MTases

| Virus | Gene | dN/dS* | Model | B_free vs B_neut model P-value | Viral DNA Methylation |
|---|---|---|---|---|---|
| Mollivirus sibericum | ml_135 | **0.044** | M0 | - | Yes |
| | ml_216 | **0.004** | b_free | <1x10$^{-6}$ | Yes |
| | ml_498 | - | - | - | No, pseudogene |
| Cedratvirus kamtchatka | ckam_cds_212 | **0.002** | b_free | 0.02 | Yes |
| Pithovirus sibericum | pv_264 | **0.055** | M0 | - | No |
| Pandoravirus salinus | psal_cds_866 | **0.281** | b_free | 0.00006 | Yes |
| | psal_cds_930 | 1 | b_neut | 0.1 | Yes |
| | psal_cds_656 | 0.354 | b_neut | 0.19 | Yes |
| Pandoravirus dulcis | pdul_cds_639 | 1 | b_neut | 0.21 | Yes |
| | pdul_cds_713 | **0.204** | b_free | 0.014 | Yes |
| | pdul_cds_889 | 0.320 | b_neut | 0.31 | Yes |
| | pdul_cds_425 | 0.658 | b_neut | 0.42 | Yes |
| Pandoravirus inopinatum | pino_cds_419 | - | - | - | Pseudogene |
| | pino_cds_741 | **0.28** | b_free | 0.0001 | - |
| | pino_cds_313 | 0.624 | b_neut | 0.66 | - |
| Pandoravirus pampulha | ppam_cds_637 | 0.493 | b_neut | 0.44 | - |
| | ppam_cds_578 | - | - | - | Pseudogene |
| | ppam_cds_790 | 0.529 | b_neut | 0.8 | - |
| Pandoravirus quercus | pqer_cds_896 | **0.242** | b_free | 0.007 | Yes |
| | pqer_cds_559 | **0.144** | M0 | - | Yes |
| Pandoravirus celtis | pcts_cds_910 | **0.361** | b_free | 0.02 | Yes |
| | pcts_cds_576 | **0.144** | M0 | - | Yes |
| Pandoravirus neocaledonia | pneo_cds_78 | **0.272** | b_free | 0.04 | Yes |
| Pandoravirus braziliansis | pbra_cds_857 | 0.357 | b_neut | 0.21 | - |
| | pbra_cds_273 | **0.243** | b_free | 0.0003 | - |
| Pandoravirus massiliensis | pmas_cds_277 | **0.164** | b_free | 0.0002 | - |
| Pandoravirus macleodensis | pmac_cds_678 | **0.304** | b_free | 0.014 | - |

*dN/dS significantly different from 1 are indicated in bold.
